# Mathematical relations between measures of brain connectivity estimated from electrophysiological recordings for Gaussian distributed data

**DOI:** 10.1101/680678

**Authors:** Guido Nolte, Edgar Galindo-Leon, Zhenghan Li, Xun Liu, Andreas K. Engel

## Abstract

A large variety of methods exist to estimate brain coupling in the frequency domain from electrophysiological data measured e.g. by EEG and MEG. Those data are to reasonable approximation, though certainly not perfectly, Gaussian distributed. This work is based on the well-known fact that for Gaussian distributed data, the cross-spectrum completely determines all statistical properties. In particular, for an infinite number of data, all normalized coupling measures at a given frequency are a function of complex coherency. However, it is largely unknown what the functional relations are. We here present those functional relations for six different measures: the weighted phase lag index, the phase lag index, the absolute value and imaginary part of the phase locking value (PLV), power envelope correlation, and power envelope correlation with correction for artifacts of volume conduction. With the exception of PLV, the final results are simple closed form formulas. We tested in simulations of linear and nonlinear dynamical systems and for empirical resting state EEG on sensor level to what extent a model, namely the respective function of coherency, can explain the observed couplings. For empirical data w e found that for measures of phase-phase coupling deviations from the model are in general minWor, while power envelope correlations systematically deviate from the model for all frequencies. For power envelope correlation with correction for artifacts of volume conduction the model cannot explain the observed couplings at all. We also analyzed power envelope correlation as a function of time and frequency in an event related experiment using a stroop reaction task and found significant event related deviations mostly in the alpha range.

## 1. Introduction

Electrophysiological recordings like electroencephalography (EEG) and magnetoencephalography (MEG) have a high temporal resolution, but are also non-invasive measurements with a low spatial resolution. The high temporal resolution allows to study brain oscillations, which are a ubiquitous phenomenon in many different frequency bands ranging from slow oscillations (around 1 Hz) to the high gamma rhythm (up to around 150 Hz). It is argued by many researchers that the functional role of these oscillations is a mechanism of communication between different brain areas [Engel et al., 2001, Fries, 2005, 2015, Engel et al., 2013]. However, it is largely unclear what features of these oscillations are relevant for which specific communication within the brain.

Oscillations at a given time point, or rather segment of time, can be characterized by the frequency, the amplitude and the phase. In principle, each of these features may serve as an independent constituent of the mechanism of the communication. It is, e.g., conceivable, that phases at two neuronal sites are strongly coupled while the amplitudes are completely independent of each other and vice versa. To study the mechanisms, measures of functional dependence have been developed which mainly focus on three kinds of coupling: phase-phase-coupling, phase-amplitude coupling and amplitude-amplitude coupling [Engel et al., 2013].

The question to be addressed here is whether the corresponding measures really describe different phenomena as it is also, at least mathematically, conceivable that, e.g., phase-phase coupling determines amplitude-amplitude coupling even if the actual values are not identical. In such a case the latter would be a function of the former; the estimation of the latter would not add information on the brain dynamics and our measures would be essentially redundant. Such a redundancy occurs if the data are Gaussian distributed. In that case linear statistics, i.e., means and cross-correlation matrices or means and cross-spectra in the Fourier domain, completely determine all statistical properties. Furthermore, all coupling measures considered in this paper are normalized and independent of global (i.e., time independent) scale transformations of the data, and then all measures must be functions of complex coherency, which is the normalized version of a cross-spectrum [Nunez et al., 1997].

In general, data are Gaussian distributed if the underlying dynamical system is linear and stationary. While, EEG and MEG data are surely not perfectly Gaussian distributed, assuming the data to be Gaussian distributed can still be a reasonable approximation. The validity of such an approximation is implicitly or explicitly assumed when estimating brain connectivity from fitting a linear dynamical model to the data as is done frequently for directed measures of connectivity like for Granger Causality [Bressler and Seth, 2011], partial directed coherence [Baccala and Sameshima, 2001], or the directed transfer function [Kaminski, 1991].

EEG and MEG have a low spatial resolution, and as consequence estimates of neuronal activities are in general mixtures of the true sources. Non-vanishing functional dependencies between such signals can be a result of such mixtures even if the underlying sources themselves are uncoupled [Nunez et al., 1997]. This is apparent on sensor level but the problem also persists on source level [Schoffelen and Gross, 2009]. To address this problem, usually referred to as ‘artifact of volume conduction’, and to remove or at least attenuate this artifact, several modifications of coupling measures were suggested exploiting the fact that the mixing is essentially instantaneous.

These two questions, what kind of coupling are we interested in and how do we remove artifacts of volume conduction, led to a large variety of coupling measures. Assuming Gaussian distributed data, all nonlinear measures must be functions of coherency, and the main content of this paper is the derivation of these functions. This allows to calculate a nonlinear coupling measure with a linear model, i.e. we can calculate from empirical data complex coherency and use the respective function as a prediction for the nonlinear measure. The difference of the two is then a measure of non-Gaussianity, and it has the potential to detect new phenomena which could otherwise be masked by linear effects.

We here analyze six of these measures plus some variations, namely four nonlinear measures of phase-phase coupling, the weighted phase lag index [Vinck et al., 2011], the phase lag index [Stam et al., 2007], the phase locking value, analyzing both the absolute value [Lachaux et al., 1999] and the imaginary part of it [Sadaghiani et al., 2012], and two measures of amplitude-amplitude coupling, one without correction for artifacts of volume conduction [Mehrkanoon et al., 2014] and one with correction for artifacts of volume conduction [Brookes et al., 2012, Hipp et al., 2012].

This paper is organized as follows. We first present background information on linear methods, i.e. coherency and basic functions of it, in section 2.1. In section 2.2 we present the procedure to find or verify mathematical relations numerically. The main part of this paper are sections 2.3 and 2.4 where we present all theoretical findings for phase-phase coupling and amplitude-amplitude coupling, respectively. We finally present results for simulations, resting state and event related EEG data in sections 3.1, 3.2 and 3.3, respectively. A conclusion is presented in section 4. We tried to keep the main body of the paper as simple as possible, and we therefore moved all mathematical derivations, which are technically quite involved, to an appendix.

## 2. Theory

### 2.1. Background on linear coupling measures

A standard approach to estimate linear relations between two electrophysiological recordings, which can be signals at sensors or estimated sources, as a function of frequency is coherency [Nunez et al., 1997]. Typically, data are divided into segments, and for each segment the data are windowed, e.g., using a Hanning window and the Fourier transformations are calculated. Alternative approaches using wavelets or the Hilbert transformation of filtered data are argued to be formally equivalent [Bruns, 2004].

The results are complex numbers *z_i_*(*f, k*) for the recordings at sensor *i* at frequency *f* and segment *k.* In the following, we will drop the frequency as argument with the implicit understanding that the analysis is done for some given frequency, and we will also omit the segment index *k* with the implicit understanding that expected values, denoted as < · >, are estimated for empirical data by averaging over *k*.

For linear and stationary dynamical systems the cross-spectrum contains complete statistical information about the system. It is defined as

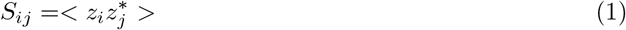

Regardless of the details of how data are defined in the frequency domain, for linear and stationary dynamical systems they are always a linear superposition of Gaussian distributed data, are hence themselves Gaussian distributed in the complex domain. Due to stationarity the distribution can only depend on phase differences and not on the phases directly. This distribution is circular Gaussian defined as [Aydore et al., 2013]

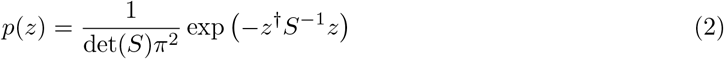

The distribution will be used below for all analytic relations between linear and nonlinear relations.

The diagonal elements of *S* are the power values, and the complex coherency *C_ij_* is calculated as

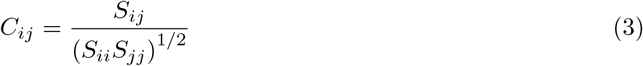

Coherency, like all other measures considered in this paper, can be calculated pairwise. To study relations between different coupling measures it is sufficient to consider only two recordings. For ease of notation, we will therefore omit the sensor index and define coherency *c* as

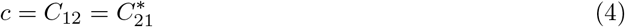

Coherency is a complex number. Its absolute value, usually called coherence, is a measure of the strength of the coupling, while its phase is a measure of the average time delay between the peaks of the oscillations. Coherency is a measure of phase-phase coupling, which, however, also depends on amplitude variations because segments of high amplitudes are weighted higher than those with lower amplitude.

The estimation of coupling using EEG and MEG sensor data, and also using respective source estimates, is prone to artifacts of volume conduction. This means that the recordings are mixtures of the true brain activities, and an estimated coupling is likely to be caused by this mixing rather than true coupling between different neuronal sites. To address this problem it was suggested to use the imaginary part of coherency, usually called ‘imaginary coherence’

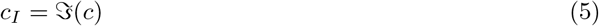

where 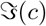 denotes imaginary part of *c*. It can be shown that *c_I_*, also denoted as ImCoh, vanishes for an infinite number of data if all brain sources are independent provided that the quasi-static approximation of the forward model is valid, i.e. the mapping of sources to sensors is instantaneous [Nolte et al., 2004]. It should be emphasized that for interacting sources the value of *c_I_*, if nonvanishing, depends on how sources are mapped into sensors.

An important measure for our analysis is lagged coherence [Pascual-Marqui, 2007, Pascual-Marqui et al., 2011], which was proposed both as a signed and unsigned version. We here use the signed version

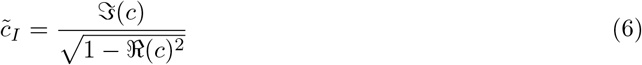

where 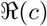 denotes real part of *c*.

Lagged coherence, also denoted as LagCoh, has the property that, in contrast to imaginary coherence, its value, apart from sign, does not depend on how the sources are mapped into sources provided that there are only two sources. Obviously, if the absolute value (or square) of 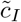 is taken, the dependence on the mapping drops out completely. However, the practical value of this might be limited, because for typical EEG or MEG measurements more than two sources (including all noise sources) are mapped into sensors. In spite of these practical limitations of its interpretation, this quantity seems to play an important and almost universal role for the relation between linear and nonlinear coupling measures as we will see below.

### 2.2. General remarks on numerical evaluation of coupling measures

The purpose of simulations is usually to demonstrate the performance of a method under realistic conditions, which we will do below in section2.2. However, our purpose is to validate mathematical relations with very high accuracy which we would like to illustrate together with theory. To verify analytical results we simulate Gaussian distributed pairs of complex numbers with a random cross-spectral matrix. For each cross-spectral matrix we use 10^7^ pairs of complex numbers. Such a simulation is not meant to represent a realistic measurement but merely to be a tool to find or verify relations with extremely high accuracy. The cross-spectra are constructed as follows. Let

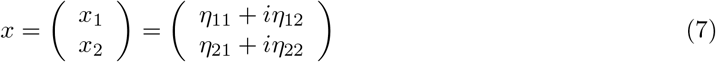

where *η_nm_* are independent Gaussian distributed real numbers with zero mean and unit standard deviation and *i* denotes imaginary unit. Then these numbers are mixed using a random complex mixing matrix *A*

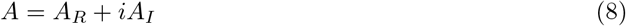

where all elements of *A_R_* and *A_I_* are independent Gaussian distributed numbers of zero mean and unit variance. For each mixing matrix A we simulate 10^7^ realizations of observations *z* as

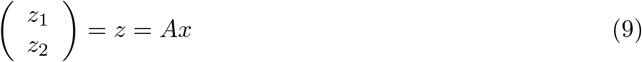

The cross-spectrum of *z* is then given by

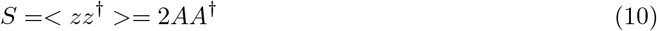

Note that *S* is in general complex because the mixing matrix *A* is complex which should not be confused with real valued mixing like a mixing artifact occurring in EEG and MEG measurements.

All coupling measures to be analyzed are constructed from expected values of the general form < *g*(*z*_1_, *z*_2_) >, with the functions *g* varying across measures, and where those expected values are estimated as averages over all realizations of *z*.

### 2.3. Phase-Phase coupling

#### 2.3.1. wPLI and PLI

Like coherency, the weighted phase lag index (wPLI) is a measure of phase-phase coupling, with averages of phase differences weighted by the amplitudes of individual segments. It is defined here as

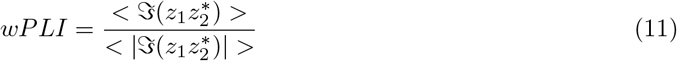

Our definition differs slightly from the original definition where in the end the absolute value is taken. Like for lagged coherence, we prefer to keep the sign because the sign might contain relevant information and in general it also simplifies statistics because the absolute value introduces a bias towards positive values. Like lagged coherence, wPLI is invariant to mixing of two sources. This is strictly true only if the absolute value is taken, but the sign may flip using our definition.

We started our tour across a series of coupling measures with wPLI because the relationship to coherency for Gaussian distributed data is known already [Ewald et al., 2012], it has a simple closed form solution, and it illustrates the typical aspects of such relations in general. The relation reads

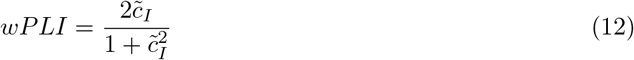

where 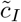 is the lagged coherence defined in Eq.6. We consider 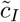 as equivalent to wPLI for Gaussian distributed data in the sense that the latter can be calculated from the former. This is exactly true only for an infinite number of data. For a finite number of data, there are, of course, statistical variations.

We emphasize two important points. First, the equivalence is not trivial, e.g. wPLI is equivalent to 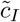 but not to imaginary coherence *c_I_*. Second, equivalent does not mean identical. The functional relation is simple here, but below we will also see other examples where we observe equivalence clearly from numerical evaluations but the precise functional relation is unclear to us. Numerical results to illustrate these findings are presented in the upper row of Fig.1.

The phase lag index (PLI) is defined as [Stam et al., 2007]

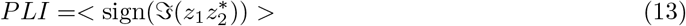

**Figure 1:**
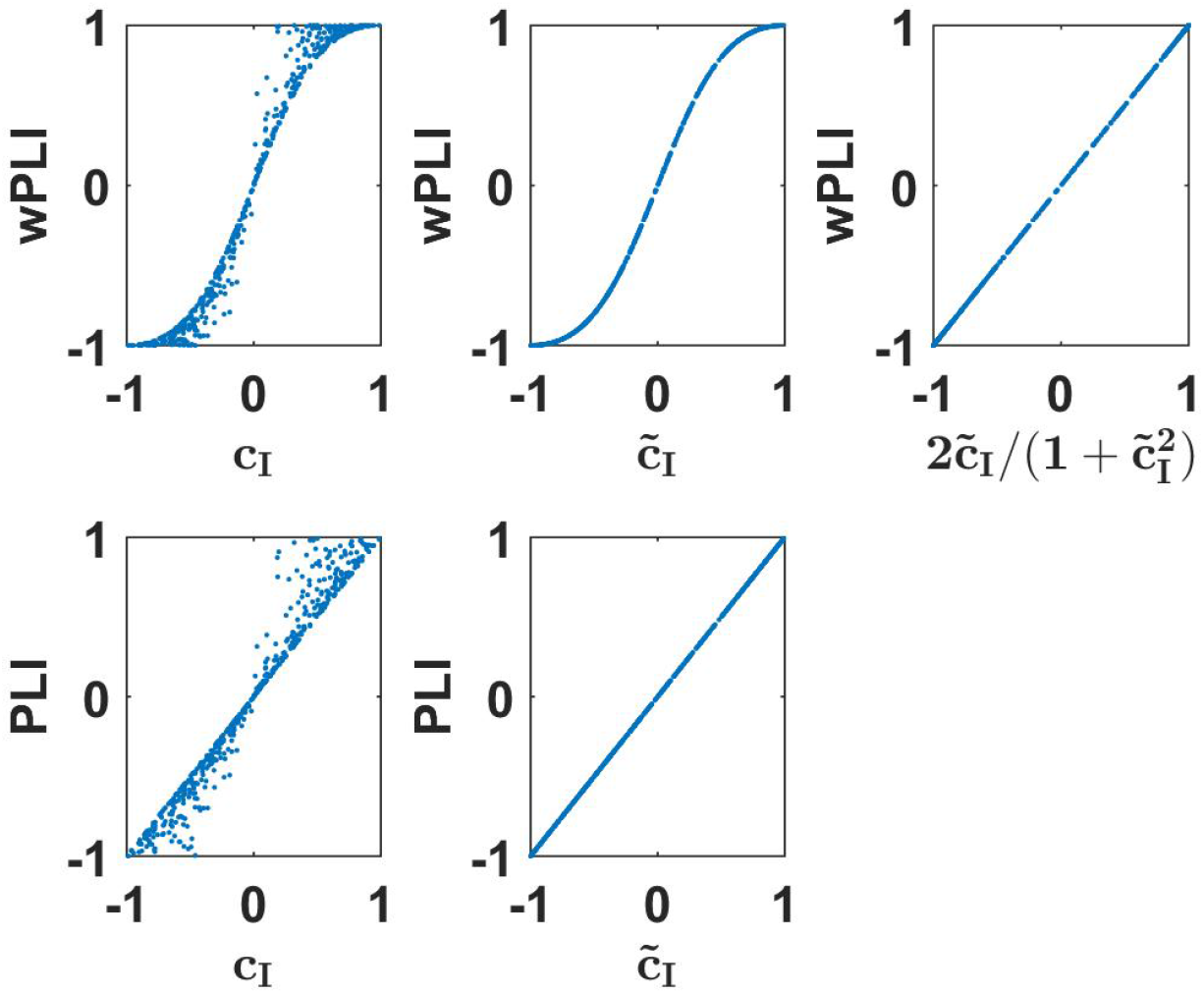
Upper row: wPLI as a function of linear measures as indicated for a 500 simulated Gaussian distributed random data sets each consisting of 10^7^ trials. Lower panels: the same for PLI.

In spite of its name, it is conceptually only loosely related to wPLI. The idea of PLI, using only the sign of the phase differences, is that the resulting coupling should be made less dependent on the actual phase difference. In practice, however, this is hardly the case because for small phase differences a sign flip is more likely than for large phase differences. In contrast to wPLI, PLI is a pure phase measure: any dependence on amplitude was removed in its definition.

As was shown by Pascual-Marqui et al. [2018], PLI, like wPLI, is invariant to mixing the signals within two sensors (or estimated sources) implying that it is invariant of the mixing of sources into sensors provided that there are not more than two sources. This can be used to analytically derive the relation between PLI and linear connectivity measures

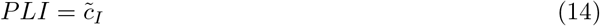

The derivation of this is presented in the appendix. Numerical results to illustrate these findings are presented in the lower row of Fig.1. We emphasize that such an identity is (apparently) true for an infinite number of Gaussian distributed data. For a finite data size, different measures have different statistical properties. The analysis of that is beyond the scope of this paper.

#### 2.3.2. PLV

The phase locking value (PLV) is a classical measure of phase-phase coupling defined *here* as a complex number:

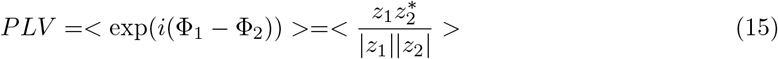

with *z_k_* = *r_k_* exp(*i*Φ_*k*_). Like PLI and in contrast to wPLI it only depends on phase differences and all amplitude variations are ignored.

This slightly deviates from the original definition, where the absolute value is taken in the end [Lachaux et al., 1999]. In Sadaghiani et al. [2012], Palva et al. [2018], Bruna et al. [2018] it was suggested to use the imaginary part of the complex definition of PLV, referred to as ImPLV in corresponding figures, to construct a measure robust to artifacts of volume conduction. We therefore prefer to keep the complex formulation and present the theory as a whole.

The analytic relation between PLV and coherency is very difficult to derive. In [Aydore et al., 2013] a solution was found in terms of hypergeometric functions using Mathematica to solve some of the integrals. In the appendix we present an explicit derivation which we found numerically to be equivalent to the solution by Aydore et al. [2013]. We get the following relation for Gaussian distributed data:

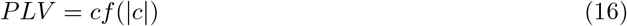

introducing a ‘scaling function’ *f* which only depends on the absolute value of coherency. We could calculate *f* analytically only as a series expansion, but not in closed form:

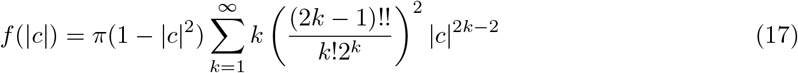

This expansion converges poorly if |*c*| is close to 1. We refer the reader to the appendix for an alternative (and less compact) formulation with better convergence properties. There we also give recommendations how to evaluate the function numerically.

We found that f is approximately linear as a function of 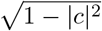, and *f* can be approximated very well by a function 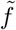 using such a linear function with exact values at the boundaries, namely *f*(0) = *π*/4 and *f*(1) = 1, leading to

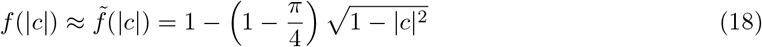

Using this approximate function, errors for PLV are smaller than .012 which we consider as negligible for practical applications. The functions *f* and 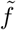 are shown in Fig.2. For all further analysis we will use the approximate scaling function 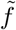.

**Figure 2:**
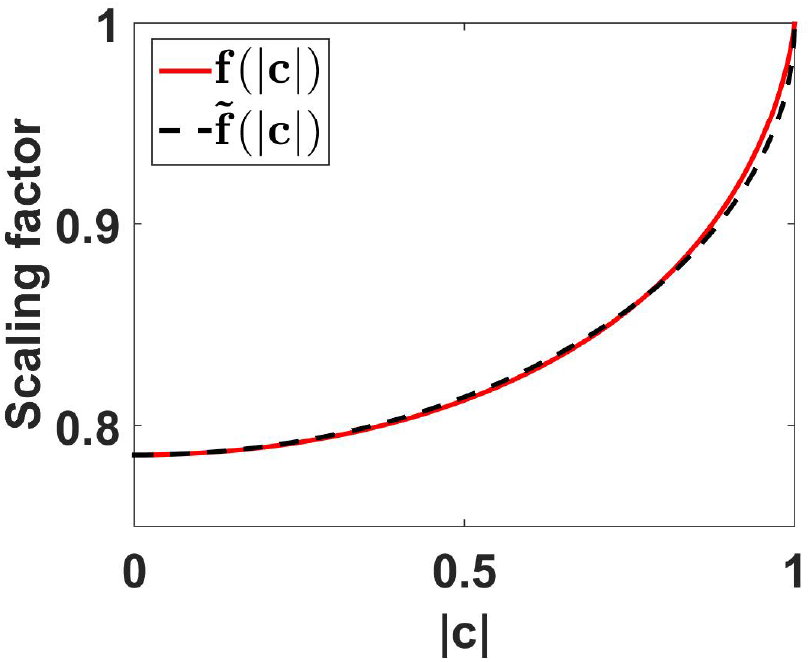
Scaling functions *f* and 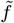 as a function of coherence. For the calculation of *f* (|*c*|) we used the first 500 terms of the expansion given in Eq.70 of the appendix.

Numerical results showing the absolute value of PLV as a function of coherence and its scaled version, and also the imaginary part of PLV as a function of imaginary coherence and its scaled version are shown in Fig.3. We observe nearly exact identities for the two scaled versions.

**Figure 3:**
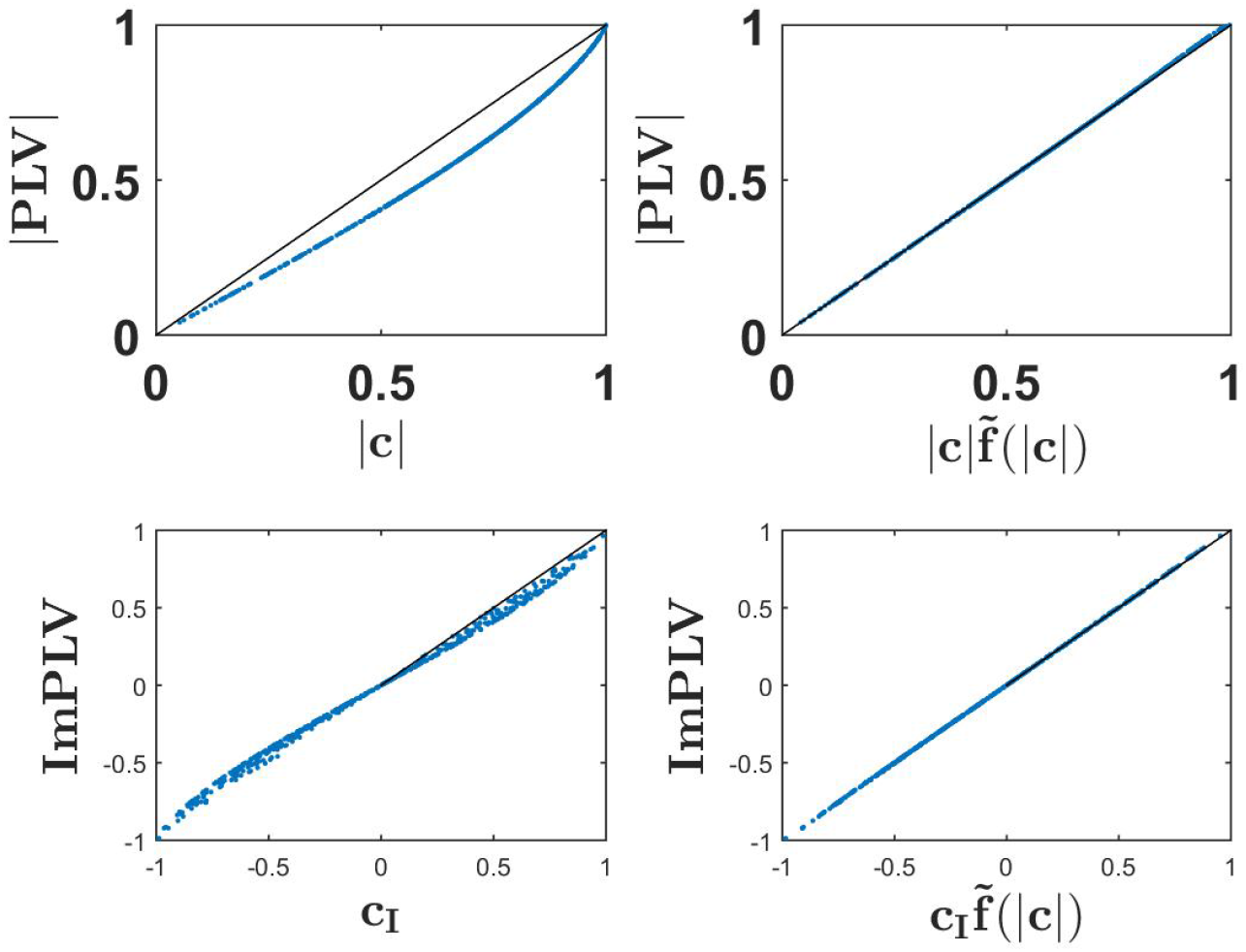
Upper row: the absolute value of PLV as a function of coherence (upper left panel) and as a function of scaled coherence (upper right)) for a 500 simulated Gaussian distributed random data sets each consisting of 10^7^ trials. Lower row: ImPLV as a function of ImCoh (left) and scaled ImCoh (right) for the same random data sets.

#### 2.4. Amplitude-Amplitude coupling

For amplitude-amplitude coupling relative phase differences are, in its original version, ignored, and the question is whether the amplitudes of two oscillation are functionally related. In principle, the two oscillations can have different frequencies. In such a case we observe cross-frequency coupling, which is always inconsistent with linear dynamical systems. Therefore, we here only consider amplitude-amplitude coupling within a specific frequency.

A functional relation between amplitudes can be measured by a correlation, but details can vary depending on what exactly is correlated. We here consider three different versions: a) the correlation between powers (the square of the amplitudes), denoted as *corr*(|*z*_1_|^2^, |*z*_2_|^2^) [Mehrkanoon et al., 2014, Soto et al., 2016], b) the correlation of amplitudes, i.e. *corr*(|*z*_1_|^2^, |*z*_2_|^2^) [Brookes et al., 2012], and c) the correlation of the logarithms of the amplitudes, i.e. *corr*(log |*z*_1_|, log |*z*_2_|) [Hipp et al., 2012]. Note, that the latter is equivalent to the correlation of the logarithm of the powers. We here use the term power envelope correlation (PEC) for all these three variants.

We have an analytic solution for Gaussian distributed data only for the first version, which reads

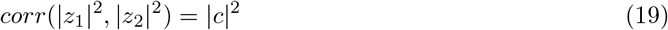

where |*c*| is the coherence of the two signals. A proof is given in the appendix. A specific consequence is that powers cannot be negatively correlated for linear dynamical systems. Note, that means are subtracted for the calculation of a correlation, and this non-negativity is not a trivial consequence of the positivity of the powers.

Like PLV and coherence, PEC is prone to artifacts of volume conduction. To address this problem it was suggested to replace *z*_2_ by *z*_2_ – *αz*_1_ where the real valued coefficient *α* is found from fitting *αz*_1_ to *z*_2_ [Brookes et al., 2012, 2014]. In the language of the original time series, this means that only that part of the second time series is evaluated which is orthogonal to the first. Without loss of generality, *z*_1_ and *z*_2_ can be normalized such that

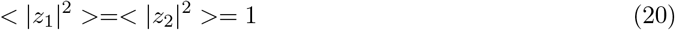

and then it is straight forward to show that

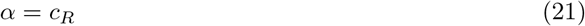

i.e. α is equal to the real part of coherency.

A different approach was proposed by Hipp et al. [2012] where it was suggested to replace |*z*_2_| by 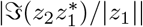. The essential difference between these two approaches is that in the first approach α is found globally, i.e., it is the same coefficient for all segments, whereas the second approach is equivalent to fitting a coefficient *α* separately for each segment. We refer here to the latter approach as a local orthogonalization, and for all of these variants we use the generic term ‘orthogonalized power envelope correlation’ (OPEC).

Similar to PEC, also for OPEC the correlation can refer to power, amplitude or logarithm of the amplitude. We have an analytic solution only for OPEC using power and for global orthogonalization. For normalized signals it reads

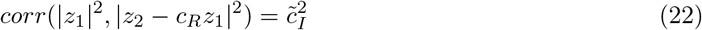

with 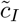 being lagged coherence given in Eq.6. The derivation of this is given in the appendix.

Numerical results for all 9 combinations (3 for PEC, 3 for OPEC with global orthogonalization, and 3 for OPEC with local orthogonalization) are shown in Fig.4. Remarkably, for all 6 OPEC variants, the coupling is apparently a unique function of lagged coherence (and then not of imaginary coherence). We also found that correlations of amplitudes are almost identical to those of powers, while the logarithmic transformation has a larger impact.

**Figure 4:**
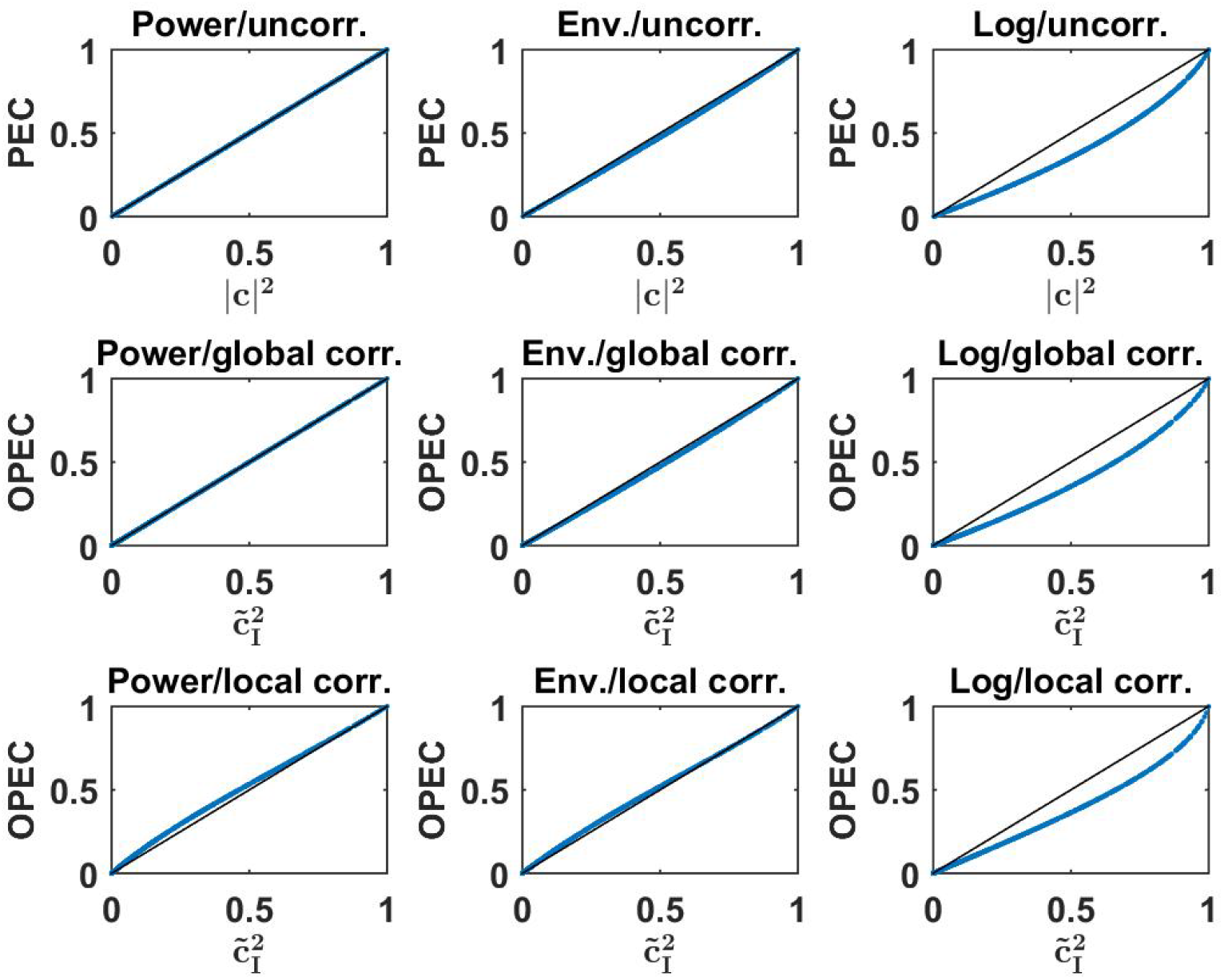
Upper row: three different versions of PEC as a function of the square of coherence for a 500 simulated Gaussian distributed random data sets each consisting of 10^7^ trials. Middle and lower row : Different versions of OPEC as indicated as a function of squared lagged coherence for the same data sets.

## 3. Results

### 3.1. Simulations

We simulated both linear and nonlinear dynamical systems. Linear dynamics was modeled with an autoregressive model as

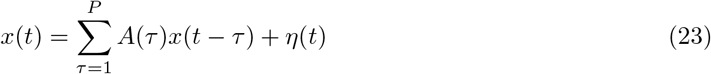

where for each discrete time point *x*(*t*) is a 2 × 1-vector for two channels, the order *P* was set to 5 and *η* was independent and white Gaussian distributed noise with zero mean and unit variance. For each data set, all coefficients of the 2 × 2 AR-matrices were set to fixed numbers randomly set from a Gaussian distribution with zero mean and a standard deviation of 0.25. Also, for each set of AR-matrices we checked analytically whether the system was stable and excluded sets with unstable dynamics. The sampling rate was understood to be 100 Hz, and, to concentrate on systematic effects, we simulated rather long data sets of 30 minutes. For a total of 100 data sets we analyzed coupling always at 10 Hz regardless of whether there was a peak in the power spectrum or not. Here and in the following data were divided onto segments of 1 second duration with 50 % overlap. The data in each segment were Hanning-windowed, and Fourier coefficients at 10 Hz were used as input for all coupling estimates.

As a nonlinear dynamical system we used the Kuramoto-model chosen here with stochastic input, random coupling parameters and random delays. Specifically, the dynamics was defined for discrete time *t* for two channel indices i as

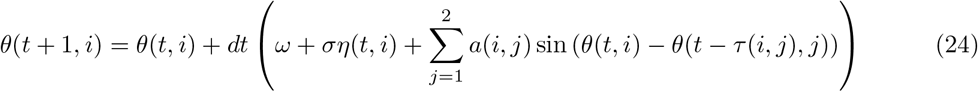

with the settings *dt* = 1/100 corresponding to a sampling rate of 100 Hz, *ω* = 20*π* resulting in 10 Hz oscillations, and *σ* = 5. The delays *τ*(*i, j*) were set at (time-independent) random integer numbers corresponding to delays up to 100 ms. The coupling parameters were set to *a*(*i, i*) = 0, and *a*(*i,j*) for *i* ≠ q was set to a random number from a Gaussian distribution with zero mean and standard deviation 0.5. Results are not crucial with respect to details of the simulations, but our choice resulted into a roughly even coverage of all possible outcomes for the linear coupling measures. From the phases the time series were finally calculated as

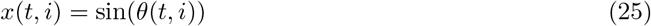

for time *t* and channel *i*. We note that even though we did not introduce an explicit time dependent amplitude, the amplitude estimate varied in time because the phases were not linear functions of time leading to variations of the instantaneous frequency and then to variations of the amplitude estimate at a specific frequency. As for the linear dynamics we simulated 100 data sets with 30 minutes of data each.

Results for both the linear and nonlinear dynamics are shown in Fig.5 where we display the nonlinear coupling value for six different coupling measures as a function of the corresponding value calculated from the linear coupling values using the derived equations for Gaussian distributed data. For the linear dynamical systems we observe nearly perfect correspondence between the two. For the nonlinear systems and for phase-phase coupling we observe only minor but clearly systematic deviations. For the nonlinear systems, amplitudes are almost not coupled at all and a prediction of that coupling assuming Gaussian distributed data fails completely.

**Figure 5:**
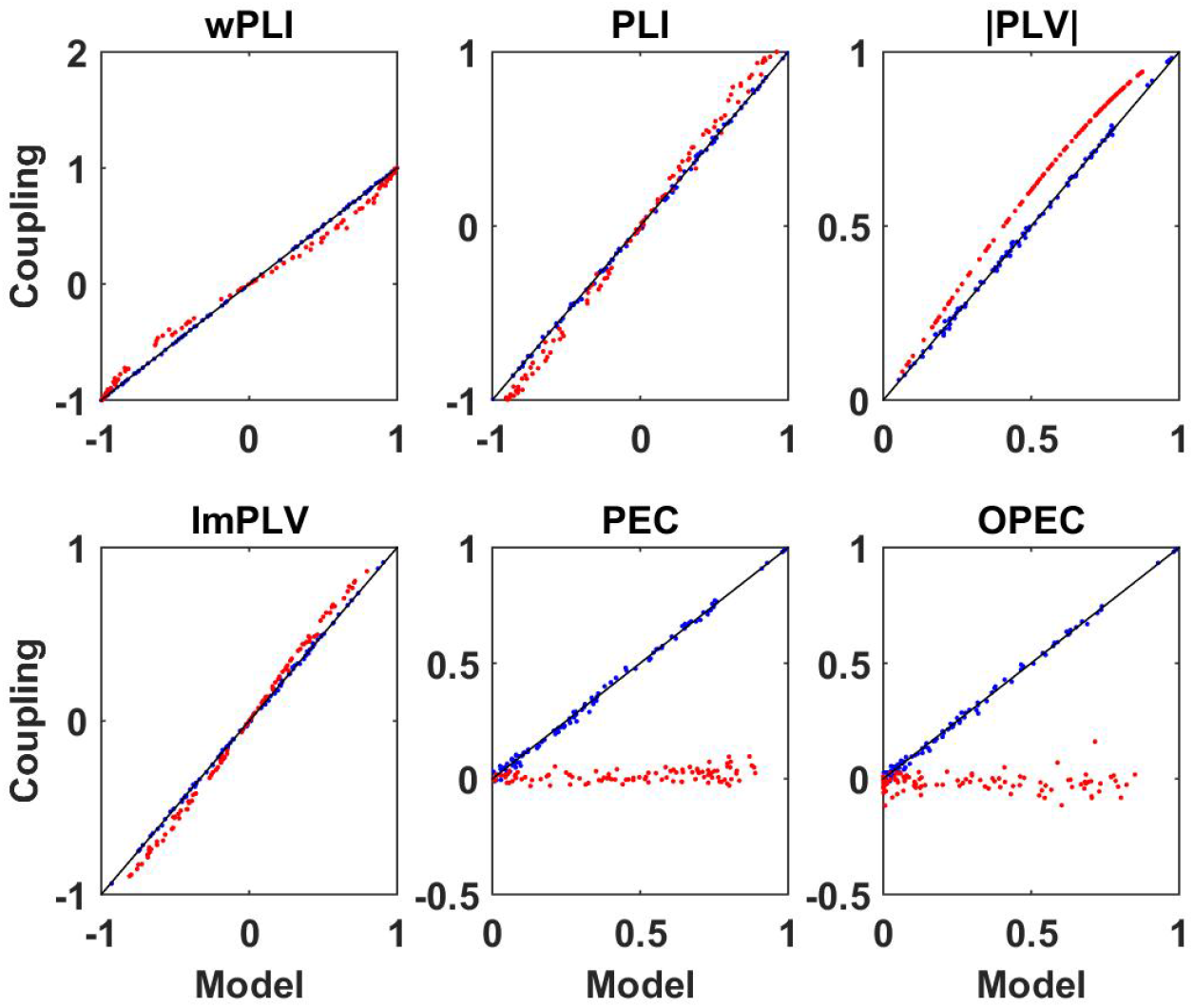
Illustration of nonlinear coupling measures as a function of their corresponding model values calculated from complex coherency and assuming Gaussian distributed data. Blue dots refer to linear dynamical systems, and red dots to the Kuramoto models each for a different specification of parameters

### 3.2. Resting state EEG

We analyzed cleaned resting state EEG data measured with eyes closed for 10 subjects publicly available at http://clopinet.com/causality/data/nolte/. The data consist of around 10 minutes recordings in 19 channels with mathematically linked ears reference. The data are used here such that our results can be reproduced. Our complete code for the analysis is available upon request. The data are a subset of data for 88 subjects, which are described in more detail in Nolte et al. [2008]. Only this subset is publicly available.

First of all, for these data sets we analyzed how well nonlinear coupling matrices can be explained by the respective linear models. Let *D*(*f, k*) be a connectivity matrix for all pairs of sensors calculated with a specific measure for frequency f and subject k, and let *D_M_* (*f,k*) be the corresponding model connectivity matrix calculated from the coherency matrix. In Fig.6 and Fig.7 we show two illustrative examples, calculated from the first subject at frequencies 10 Hz and 15 Hz, respectively. For each pair of sensors we show the actual nonlinear measure and the result of the corresponding model. Resting state with eyes closed is known to show a strong 10 Hz rhythm (the alpha rhythm) consisting of interacting sources with substantial phase delays. Such interactions are also observable at other frequencies, but typically to a much lesser extent.

**Figure 6:**
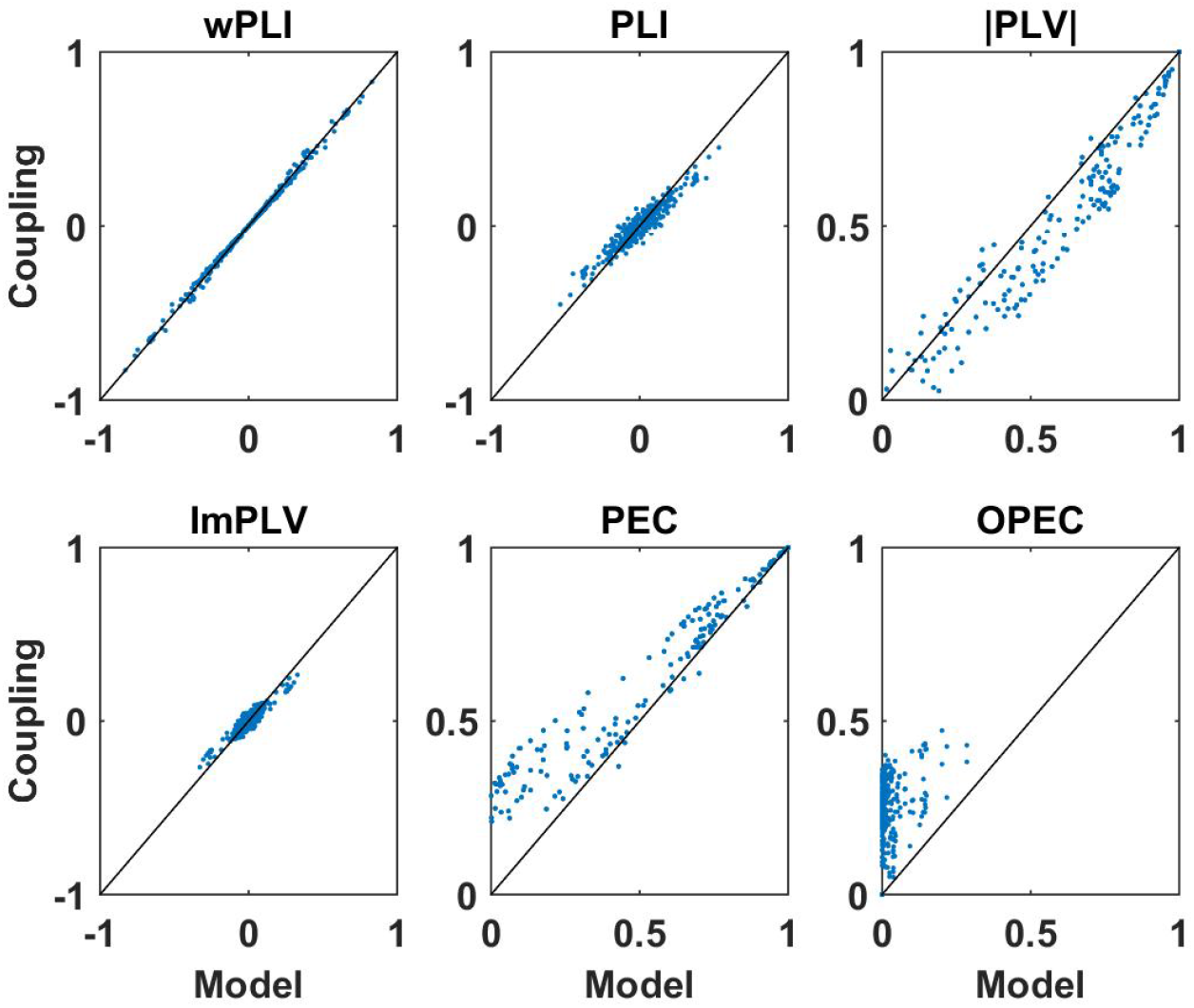
Illustration of nonlinear coupling measures as a function of their corresponding model values calculated from complex coherency and assuming Gaussian distributed data. Each dot refers to one pair of sensors for one subject with coupling values calculated at 10 Hz. For PEC and OPEC, the nonlinear measures can also come out negative, but that’s vary rare, values are typically only slightly below zero, and such a case did not occur in this example.

**Figure 7:**
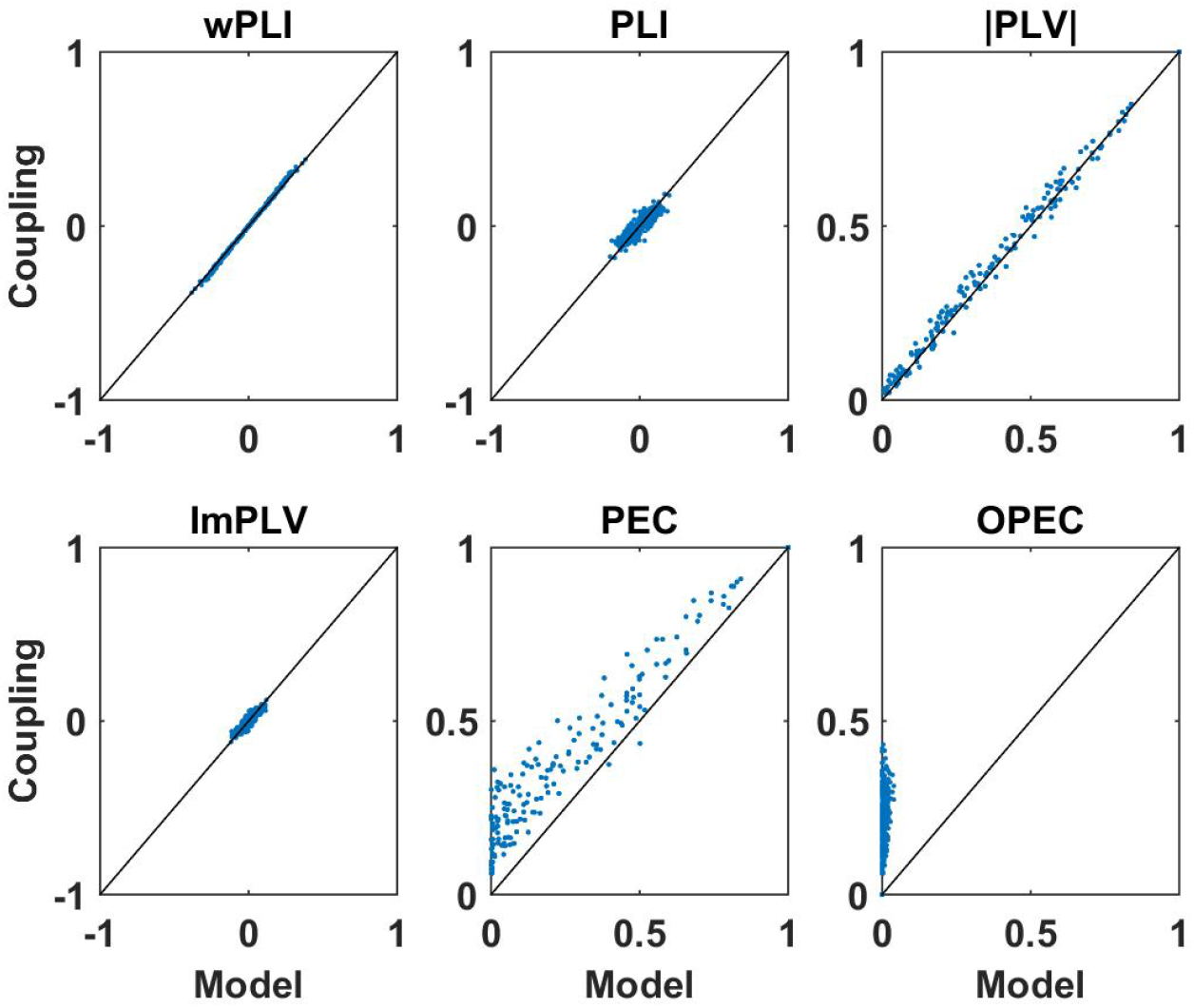
Same as Fig.6 but now for coupling measures calculated at 15 Hz. We note again that PEC and OPEC can be negative, but such a case did not occur in this example.

We consider the following observations as most relevant. The wPLI can be explained by the model almost perfectly, regardless of whether there are substantial interactions (with delay) or not, amplitude-amplitude coupling (i.e., PEC) is systematically larger than the model prediction, and amplitude-amplitude coupling with attenuation of artifacts of volume conduction (i.e., OPEC) cannot be explained by the model at all. This could be a problem when interpreting OPEC as a coupling measure robust to artifacts of volume conduction as will be discussed in the conclusion in section 4.

In the following we present a systematic analysis including results for all subjects and frequencies. For *K* subjects we define an average model error for each frequency as

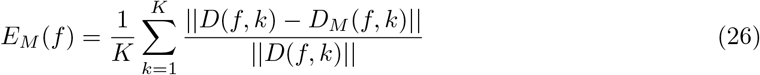

where ||·|| denotes Frobenius norm.

Non-vanishing model errors can have two causes: a) the data are non-Gaussian distributed, and b), they are caused by statistical fluctuations. To assess the magnitude of the statistical fluctuation we also calculated a statistical error. For this we replaced the model connectivity by a connectivity matrix calculated with the nonlinear measure with resampled data using the bootstrap procedure where we constructed new data of the same size by randomly picking segments of the original data with repetition. For the *k.th* subject and frequency *f* we calculated *N* = 20 such connectivity matrices denoted as *D_S_*(*f, k, n*) for *n* = 1..*N*, and a statistical error was calculated as

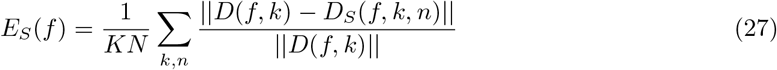

Results for 6 different nonlinear coupling measures are shown in Fig.8. In addition, we also calculated the statistical errors for the linear measures. We observe that generally for phase-phase coupling all methods with correction for artifacts of conduction are similar: results are statistically unstable for frequencies outside the alpha band. For the frequencies outside the alpha band the models are typically poor which is probably not surprising as these effects are relatively weak and can hardly be reproduced with different methods. An exception is wPLI which can always be explained very well with the linear model, indicating that wPLI depends very little on nonlinear properties of the data. The absolute value of PLV can typically be explained very well with the linear model, but to lesser extent in the alpha band. This is the only phenomenon for phase-phase coupling, where we can clearly observe deviations of the model and actual coupling larger than statistical error.

**Figure 8:**
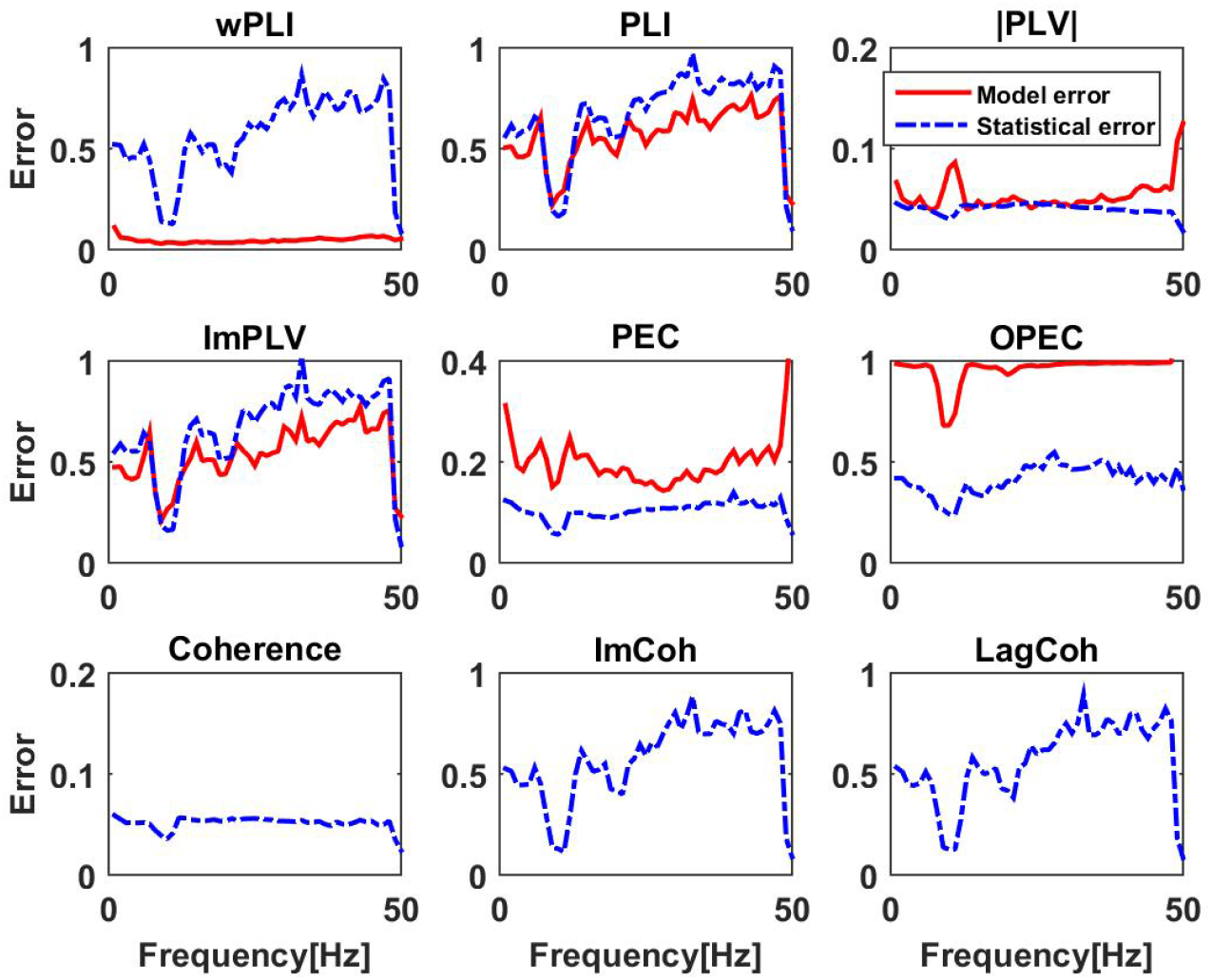
Upper and middle row: relative model errors (full lines) averaged over 10 subjects for 6 different nonlinear coupling measures each of them modeled by the corresponding function of coherency assuming Gaussian distributed data. Estimated statistical errors are shown as dashed lines. Lower row: statistical errors for three different linear measures.

Systematic deviations of the model predictions larger than statistical errors can be observed for PEC and OPEC for all frequencies. While for PEC the model (which is the square of coherence) explains around 80% of the observation (model error is around .2), the model error for OPEC is nearly 100% which is not surprising when inspecting the examples shown in Fig.6 and Fig.7.

### 3.3. Reaction task with and without conflict

We now present results for an event related paradigm. The purpose here is merely an illustration to show that we can get non-trivial results also in such a case. Details of the experiment can be found in Li et al. [2015], and we here only give a short description. In this paradigm, the stimulus was an upward or downward arrow presented at one of four possible locations of the screen: top left, top right, bottom left, and bottom right. Participants were asked to respond to the direction of the arrow as soon as possible by pressing the “F” key or the “J” key on a keyboard, while ignoring the location of the arrow, which was either congruent or incongruent with the direction of the arrow. The mapping of arrow direction and response key was counterbalanced between participants. EEG was measured in 62 channels (plus 2 mastoids, which were not included in the connectivity analysis) and referenced to the mathematically linked mastoids. In total, we analyzed 33 subjects with an average of 228 trials per condition. For each trial, 1200 ms of data from 200 ms before the stimulus until 1 second after the stimulus were analyzed further. ERPs were subtracted from the raw data such that a connectivity analysis corresponds to the analysis of fluctuations around the ERPs. Each trial was divided into segments of 200 ms duration with an overlap of 180 ms such that we could calculate connectivity for 51 different time points. Of course, such short segments of 200 ms result in a poor frequency resolution of Δ*f* ≈ 5 Hz, but with a high frequency resolution time dependence cannot be analyzed anymore.

For these data, we only analyzed PEC and calculated the correlations of the squares of the amplitudes. We recall that the model assuming Gaussian distributed data predicts that PEC is the square of coherence. Similar to the analysis of the resting state data we calculated a model error with Eq.26, which now also depends on the time of the segments relative to the stimulus. We observed that results are very similar for conflict and non-conflict trials (not shown), and we therefore present only results where we combined the conditions. The model error is shown in the upper left panel of Fig.9. The model error is relatively large (around .2) for the alpha range before the stimulus and at the end of trials. It drops substantially in the center of the trials.

**Figure 9:**
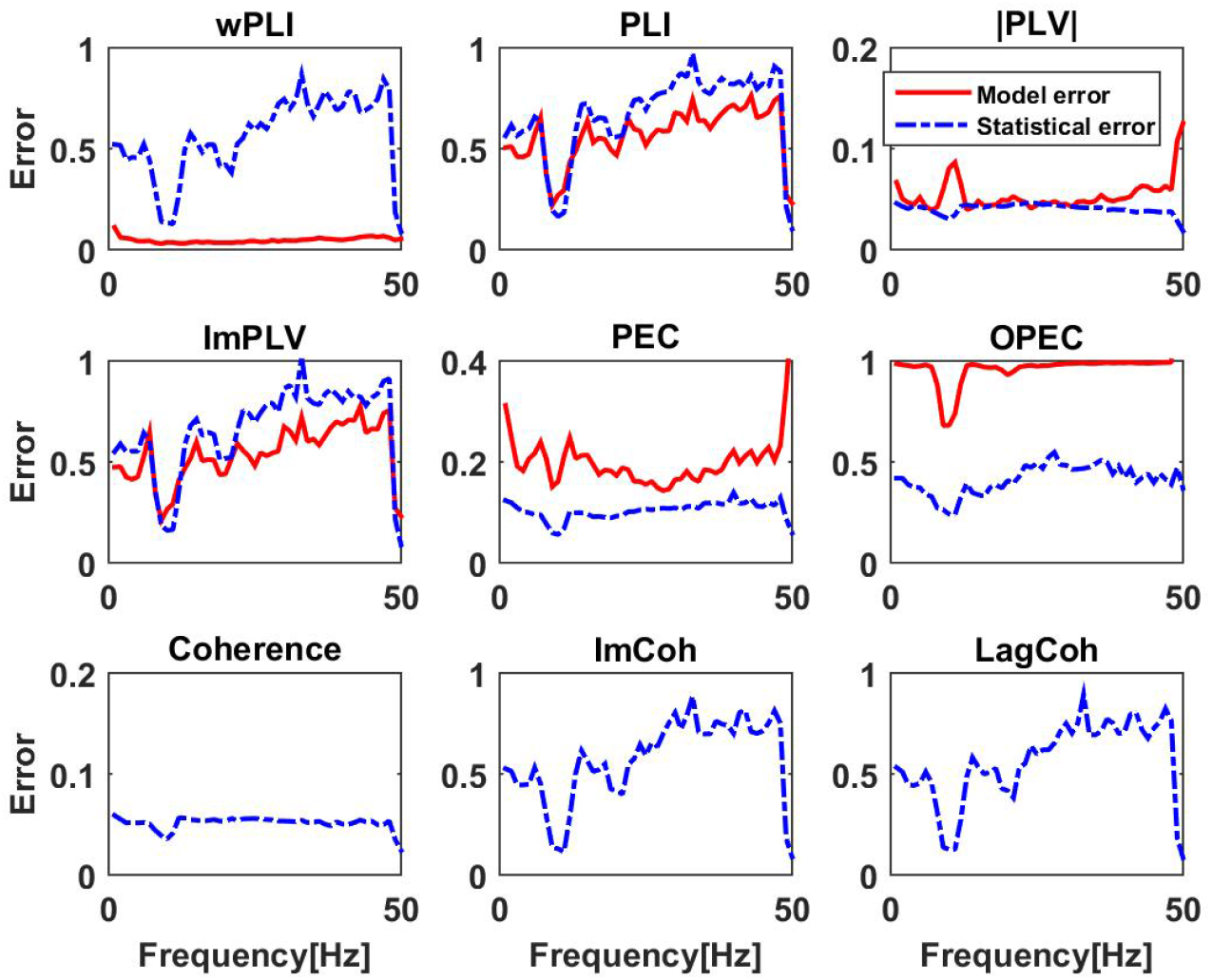
Upper row: model errors as a function of time and frequency for PEC using correlation of powers. The stimulus was at *t* = 0*ms*. In the left panel we show original model errors, and in the right panel we show model errors with results at baseline (the first 200 ms) subtracted, and with non-significant values set to zero. Lower panels: the same for powers, but here we always normalize power values by the power values at the baseline.

The question here is whether changes in time can be detected significantly. Therefor, we calculated the difference of the model error to the baseline, which we set to be the results at the first time point, i.e. the segments before the stimulus. Significance was tested using a paired permutation test: for each subject and for each time point, results (at the same frequency) were randomly switched between the baseline and the actual time point. We constructed 10.000 such surrogates and the p-value was calculated as the fraction of cases for which the surrogates showed a larger difference of baseline and actual time point than the original data. We analyzed 9 frequencies from 5 Hz to 45 Hz. Using the Bonferroni correction for 51 time points and these 9 frequencies we considered results as significant if the p-value was lower than .05/(51 * 9). In the upper right panel of Fig.9 we show the model error with baseline subtracted and after setting non-significant differences to zero. We observe significant time variation of the model error mainly in the alpha range.

In the lower panels of Fig.9 we show analogous results for power, here always showing the power ratio of the power for each time point and the corresponding power at baseline. While we observe a drop in alpha power (lower left panel), this drop is not significant.

## 4. Conclusion

In this paper we presented mathematical relations between linear and nonlinear measures of brain coupling assuming Gaussian distributed data. All relations were verified in simulations. Let us recall the main theoretical results. We considered four different nonlinear measures of phase-phase coupling: wPLI, PLI, and absolute value and imaginary part of PLV. The functional relations could be proven for all of these measures. For amplitude-amplitude coupling we could solve the problem analytically in closed form only if powers (rather than amplitudes or the logarithm of powers) are correlated and if artifacts of volume conduction are corrected for globally (i.e., time independent) or not at all. All other variants could only be analyzed numerically. To our own surprise we found that all considered versions of amplitude-amplitude coupling with correction for mixing artifacts turned out to be functions of lagged coherence. Except for the one case we could solve analytically, we must leave the mathematical proof of this as an open question. In total, with the exception of the imaginary part of PLV, all considered measures with correction for mixing turned out to be functions of lagged coherence for Gaussian distributed data, and hence lagged coherence plays a quite universal role.

In general, calculating linear measures from the cross-spectrum has a couple of computational advantages. First, if the data are Gaussian distributed, then the cross-spectrum is a maximum likelihood estimator of the parameters of the distribution and then the estimation has the smallest statistical error. Second, when using a linear inverse method (or a quasi-linear method like a beamformer) the cross-spectrum in source space can easily be calculated from the cross-spectrum in sensor space by multiplying the latter with the spatial filter from the left and right. In contrast, when using a nonlinear method, the entire raw data need to be mapped into source space. Third, in case of amplitude-amplitude coupling phase information contained in complex coherency gets lost. The question then is whether for empirical data the calculation of nonlinear measures in particular in source space is worth the effort and adds any useful information. Empirical electrophysiological data are certainly not Gaussian distributed, even if the deviation of Gaussianity is usually weak. The mathematical relations presented here may serve as a tool to isolate nonlinear effects by subtracting from the nonlinear measure the corresponding model results assuming a Gaussian distribution. Such a difference cannot be explained by any linear dynamical model. If, e.g., a substantial non-vanishing difference is observed, this could indicate that a linear autoregressive model is inadequate to calculate Granger Causality.

For empirical data we analyzed how well observed nonlinear coupling measures could be explained by calculating coherency from these data and then predicting the nonlinear coupling measure assuming Gaussian distributed data and using the theoretical relations. For event related data, we only illustrated the procedure and could show that deviations of the model for amplitude-amplitude coupling are in general time dependent. A more complete analysis was given for resting state EEG data. For phase-phase coupling we found that deviations are minor, i.e., essentially within statistical errors, with the exception of the absolute value of PLV at 10 Hz. Deviations were quite substantial for amplitude-amplitude coupling, in particular when a correction for artifacts of volume conduction is included. Since that measure is known to be robust to artifacts of volume conduction in general only for Gaussian distributed data, as made clear in much detail by the authors themselves [Brookes et al., 2014], the question remains open whether the large deviations from the Gaussian model are artifacts of volume conduction or correspond to genuine nonlinear brain interactions.

This paper leaves many questions open. Apart from the lack of mathematical proofs for some cases, the analysis of empirical data was rather coarse. We estimated model errors as averages over all sensor pairs. Interesting effects, e.g., delayed brain interactions at other frequencies than 10 Hz, certainly exist but can easily be masked by such averages if those effects are relatively weak and/or occur only in a few sensor pairs. Also, our analysis was done completely in sensor space, and the question remains open where in the brain we observe large or small deviations from the linear model, and, most importantly, whether those differences can be explained by a reasonable model of brain dynamics. Finally, the question is whether we can observe differences of these deviations for different experimental conditions or brain pathologies. All these questions are beyond the scope of this paper and need to be addressed in the future.

## Acknowledgment

This research was partially funded by the BMBF (161A130), the German Research Foundation (DFG, SFB936/A2/A3/Z3 and TRR169/B1/B4 and SPP2041/EN533/15-1), and from the Landes-forschungsförderung Hamburg (CROSS, FV25).

## Appendix

### General remarks

Here we derive analytic results for the relation between linear and nonlinear coupling measures assuming Gaussian distributed data. The order of the measures in this appendix is determined by mathematical complexity and differs from the order in the main body of this paper. All considered measures are bivariate, and, being normalized quantities, also independent of global scales. Without loss of generality we can therefore set the cross-spectrum to be

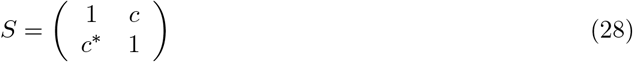

where *c* is the complex coherency. The stochastic variable is the vector consisting of two complex number *z* = (*z*_1_ *z*_2_)^*T*^ and for the following analysis we assume a Gaussian distribution with probability density function:

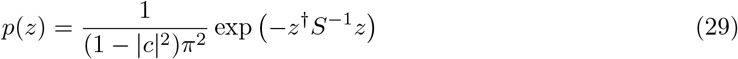

Coupling measures are in general constructed from expected values of functions *g*(*z*)

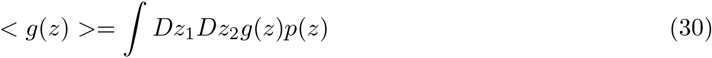

In generic form, *Dz* denotes the infinitesimal element for the integration over a complex plain. For all our analysis we use spherical coordinates, *z* = *r* exp(*i*Φ), and then *Dz* = *rdrd*Φ.

The integrals to be evaluated below will have two different forms. First, integrals of phases will always be of the form

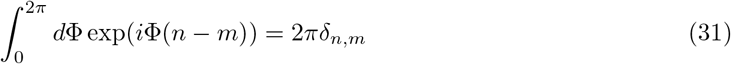

where *δ_n,m_* denotes the Kronecker delta function.

Second, integrals over amplitudes r will always be integrals of a polynomial multiplied with a Gaussian function. Those integrals are standard and read for integer *n* and *k*

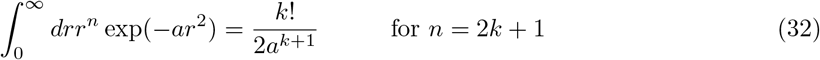

and

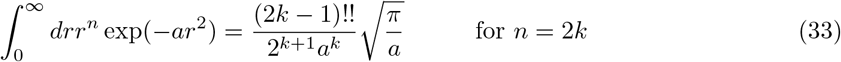

where (2*k* – 1)!! denotes the product of all odd integers from 1 to 2*k* – 1.

### Power envelope correlation

### PEC without suppression of mixing artifacts

Power envelope correlation (PEC) between two complex variables *z*_1_ and *z*_2_ is defined as the usual correlation calculated for the powers |*z*_1_|^2^ and |*z*_1_|^2^

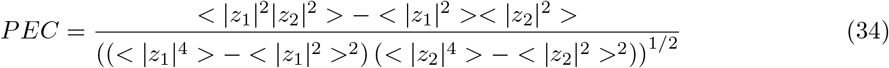

All expected values are to be evaluated for low order polynomials of *z*_1_ and *z*_2_. This can be solved in closed form using a coordinate transformation. Also, power-power correlation is independent of the phase of coherency and we can therefore assume without loss of generality that c is real valued. Let

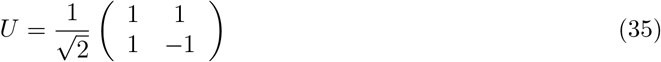

*U* is real-valued, symmetric and orthogonal, i.e. *U* = *U^T^* = *U*^†^ = *U*^−1^. This *U* diagonalizes *S* and *S*^−1^, specifically

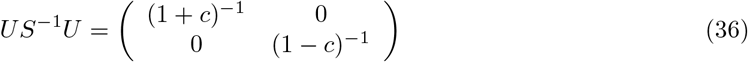

We now define new coordinates as

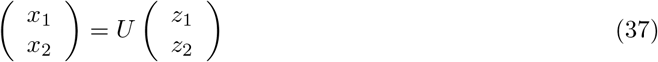

Since, det(*U*) = 1 we have for the infinitesimal elements *D*_*x*1_*D*_*x*2_ = *D*_*z*1_*D*_*z*2_. The exponent of the exponential function reads apart from the overall sign

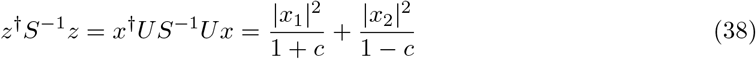

Also, the polynomials as functions of *z*_1_ and *z*_2_ occurring in Eq.34 can be expressed as polynomials of the new coordinates, e.g.

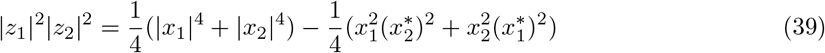

Note, that in the first two terms the dependence on the phases drops out, which is not the case for the second two terms. These second two terms do not contribute to the expected value because these contributions vanish after integration with respect to the phases according to Eq.31. The expected values of the first two terms can be directly evaluated:

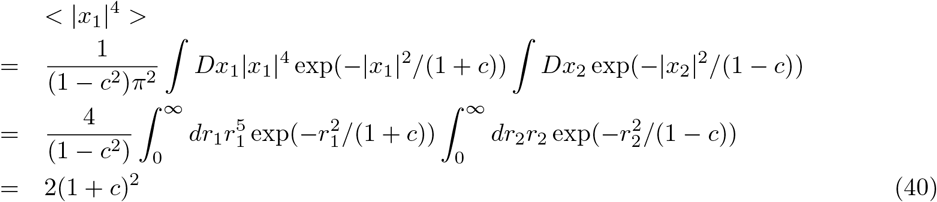

Likewise

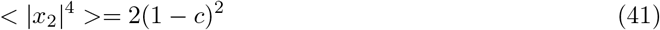

These terms can be combined to give

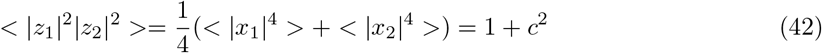

All other terms do not depend on *c* and we just present the results

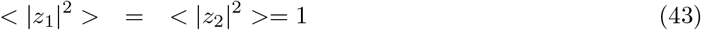

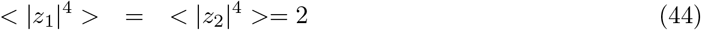

Inserting all these results into Eq.34 leads for real valued *c* to

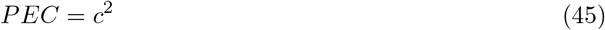

and in general, since PEC does not depend on the phase of *c*, to

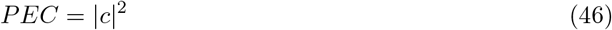

### PEC with suppression of mixing artifacts

PEC with suppression of mixing artifacts depends on phase differences, and the simplification to assume with loss of generality real valued coherency is not possible. However, the strategy to solve this problem is the same as in the previous section. We will present here only the main principles.

We want to calculate the correlation of |*z*_1_|^2^ and |*z*_2_ – *c*_*R*_*z*_1_|^2^, i.e.

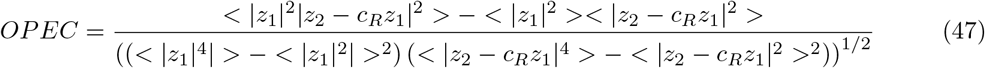

for coherency *c* = *c_R_* + *ic_I_* = |*c*| exp(*i*Φ).

Now, let

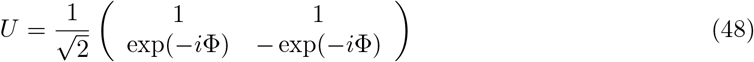

*U* is unitary, i.e. *U*^†^ = *U*^−1^, and it diagonalizes *S*^−1^

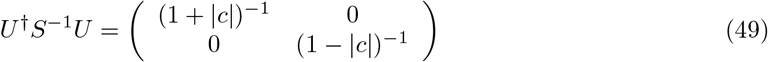

Using new coordinates *x* = *U*^†^*z* we get for the exponent in the probability density apart from the overall sign

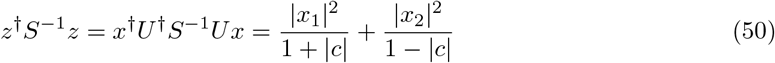

As before, all functions of z are now expressed as functions of *x* according to 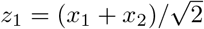 and 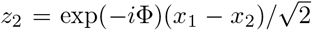. All terms are then products of functions of either *x*_1_ or *x*_2_. Integrals over phases vanish unless the respective term is not phase dependent, and the integrals over the radial variables reduce to one-dimensional integrals which can all be calculated with Eq.32. These calculations are still very tedious and we here report only the results for the individual terms. In addition to Eq.43 and Eq.44 we get

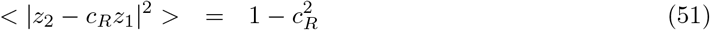

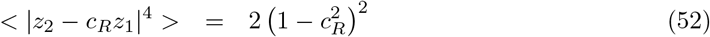

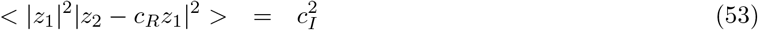

Inserting these results into Eq.47 we arrive at the final solution

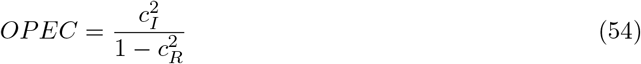

### Phase Lag Index

The calculation for PLI is similar to the one for PEC, but the integrals are slightly more complicated. We first recall that PLI is invariant to real valued linear transformations. This can be exploited to consider the case of purely imaginary coherence. If *S* and *U* are given as in Eq.28 and Eq.35, respectively, then with *c* = *c_R_* + *ic_I_* we get

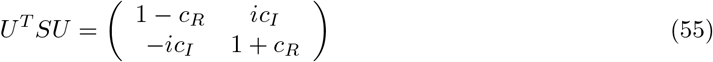

Scaling to unit diagonal elements with

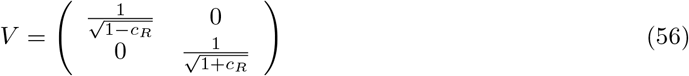

leads to

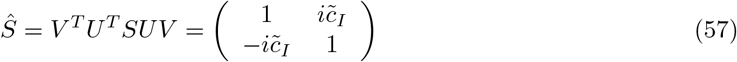

with

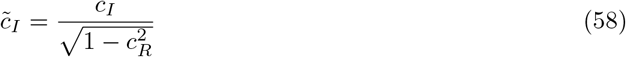

Since PLI is invariant to the above transformations, we observe that it must be a function of lagged coherence 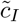. To calculate the functional form we need another transform which actually diagonalizes *S*. This can be achieved with

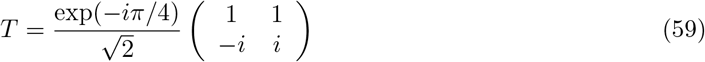

for which det(*T*) = 1 holds. Denoting the original coordinates (corresponding to covariance matrix *Ŝ*) as *z*, and defining new coordinates as *x* = *T*^†^*z* we get

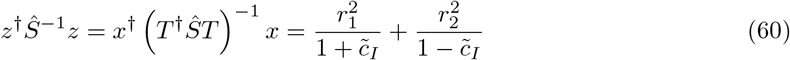

with *r_i_* = |*x_i_*|, and

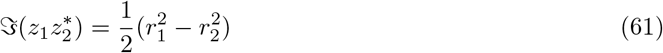

Note that phase dependencies disappear in the new coordinates and in the following we only sketch the major steps of the rather tedious but straight forward integrations along radial coordinates:

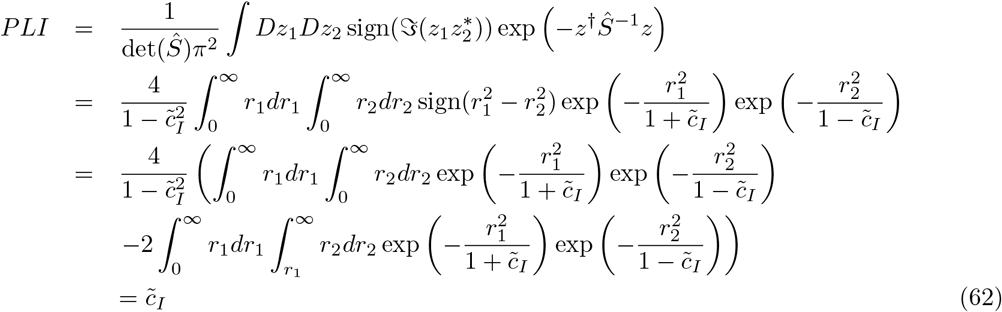

### Phase locking value

The complex phase locking value (PLV) is defined as

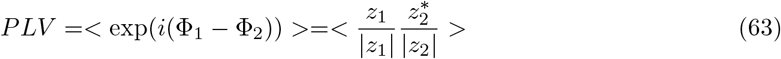

In contrast to power-power correlation, this coupling measure cannot be expressed in terms of expected values of low order polynomials of *z*_1_ and *z*_2_. As a consequence, a coordinate transformation, as was done for PEC, is not useful here. For PLV we will therefore follow a different strategy. Since we here derive relations for the full complex PLV we cannot restrict ourselves to real valued c. We rewrite the exponential function using the abbreviation *b* = 1 – |*c*|^2^

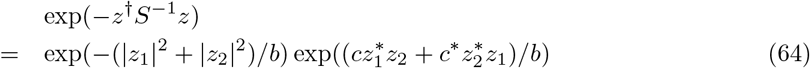

The second exponential function is expanded in a Taylor series

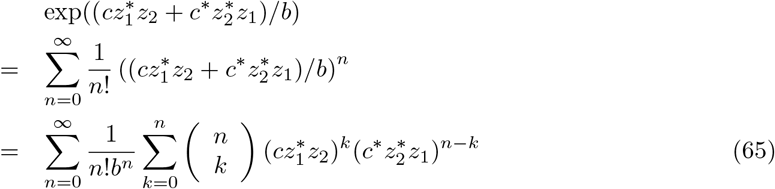

The double sum over *n* and *k* will reduce to a single sum over *n* after integration with respect to the phases. We recall that *z_k_* = *r_k_* exp(*i*Φ_*k*_). Apart from factors, which do not depend on phases, and including the factor 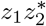 from the measure itself, we get for the integrals

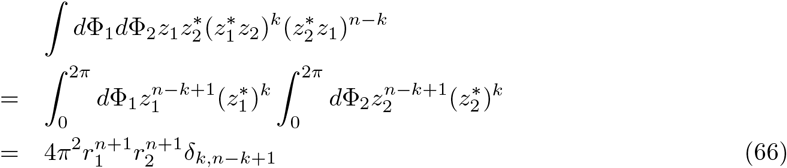

All remaining integrals are products of two one-dimensional integrals and can be evaluated with Eq.33 to give

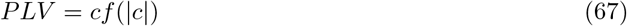

with a correction factor

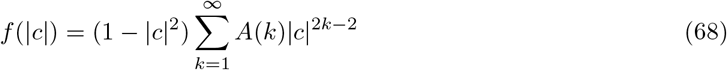

with coefficients

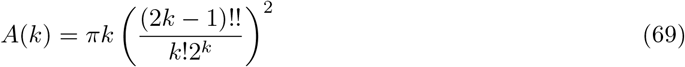

We recall that (2*k* – 1)!! denotes the product of all odd integers from 1 to 2*k* – 1 .

The series expansion of Eq.68 converges poorly if |c| is close to 1. We therefore recommend an equivalent formulation as

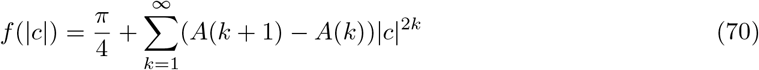

We also recommend that for the calculation of (2*k* – 1)!!/(*k*!2^*k*^) numerator and denominator should not be evaluated separately, but the whole ratio should rather be calculated as

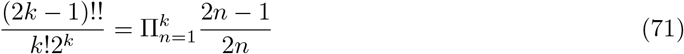

## References

Aydore, S., D. Pantazis, and R. Leahy (2013). A note on the phase locking value and its properties. Neuroimage 74, 231–244

Baccala, L. and Sameshima, K. (2001). Partial directed coherence: a new concept in neural structure determination. Biol. Cybern. 84, 463–474

Bressler, S. L. and Seth, A. K. (2011). Wiener-Granger Causality: A well established methodology. Neuroimage 58, 323–329. doi:10.1016/j.neuroimage.2010.02.059

Brookes, M. J., O’Neill, G. C., Hall, E. L., Woolrich, M. W., Baker, A., Corner, S. P., et al. (2014). Measuring temporal, spectral and spatial changes in electrophysiological brain network connectivity. Neuroimage 91, 282–299. doi:10.1016/j.neuroimage.2013.12.066

Brookes, M. J., Woolrich, M. W., and Barnes, G. R. (2012). Measuring functional connectivity in MEG: A multivariate approach insensitive to linear source leakage. Neuroimage 63, 910–920. doi: 10.1016/j.neuroimage.2012.03.048

Bruna, R., Maestu, F., and Pereda, E. (2018). Phase locking value revisited: teaching new tricks to an old dog. J. Neural Engin. 15. doi:10.1088/1741-2552/aacfe4

Bruns, A. (2004). Fourier-, hilbert- and wavelet-based signal analysis: are they really different approaches? J Neurosci Methods 137, 321–32

Engel, A. K., Fries, P., and Singer, W. (2001). Dynamic predictions: Oscillations and synchrony in top-down processing. Nature Reviews Neuroscience 2, 704–716

Engel, A. K., Gerloff, C., Hilgetag, C. C., and Nolte, G. (2013). Intrinsic Coupling Modes: Multiscale Interactions in Ongoing Brain Activity. Neuron 80, 867–886. doi:10.1016/j.neuron.2013.09.038

Ewald, A., Marzetti, L., Zappasodi, F., Meinecke, F. C., and Nolte, G. (2012). Estimating true brain connectivity from EEG/MEG data invariant to linear and static transformations in sensor space. Neuroimage 60, 476–488. doi:10.1016/j.neuroimage.2011.11.084

Fries, P. (2005). A mechanism for cognitive dynamics: neuronal communication through neuronal coherence. Trends Cogn Sci 9, 474–480

Fries, P. (2015). Rhythms for cognition: Communication through coherence. Neuron 88, 220–235. doi:10.1016/j.neuron.2015.09.034

Hipp, J. F., Hawellek, D. J., Corbetta, M., Siegel, M., and Engel, A. K. (2012). Large-scale cortical correlation structure of spontaneous oscillatory activity. Nat Neurosci 15, 884. doi:10.1038/nn.3101

Kaminski, K., MJ Blinowska (1991). A new method of the description of the information-flow in the brain structures. Biol Cybern, 203–210. doi:10.1007/BF00198091

Lachaux, J., Rodriguez, E., Martinerie, J., and Varela, F. (1999). Measuring phase synchrony in brain signals. Hum Brain Mapp 8, 194–208. doi:10.1002/(SICI)1097-0193(1999)8:4:194::AID-HBM4¿3.0.CO;2-C

Li, Q., Wang, K., Nan, W., Zheng, Y., Wu, H., Wang, H., et al. (2015). Electrophysiological dynamics reveal distinct processing of stimulus-stimulus and stimulus-response conflicts. Psychophysiology 52, 562–571

Mehrkanoon, S., Breakspear, M., Britz, J., and Boonstra, T. (2014). Intrinsic coupling modes in source-reconstructed electroencephalography. Brain Connectivity 10, 812–25

Nolte, G., Bai, O., Wheaton, L., Mari, Z., Vorbach, S., and Hallett, M. (2004). Identifying true brain interaction from eeg data using the imaginary part of coherency. Clinical Neurophysiology 115, 2292 – 2307. doi:DOI:10.1016/j.clinph.2004.04.029

Nolte, G., Ziehe, A., Nikulin, V., Schlögl, A., Krämer, N., Brismar, T., et al. (2008). Robustly estimating the flow direction of information in complex physical systems. Phys Rev Lett 100, 234101

Nunez, P., Srinivasan, R., Westdorf, A., Wijesinghe, R., Tucker, D., Silberstein, R., et al. (1997). EEG coherency. 1. Statistics, reference electrode, volume conduction, Laplacians, cortical imaging, and interpretation at multiple scales. Electroencephalogr. Clin. Neurophysiol. 103, 499–515

Palva, J. M., Wang, S. H., Palva, S., Zhigalov, A., Monto, S., Brookes, M. J., et al. (2018). Ghost interactions in MEG/EEG source space: A note of caution on inter-areal coupling measures. Neuroimage 173, 632–643. doi:10.1016/j.neuroimage.2018.02.032

Pascual-Marqui, R. D. (2007). Instantaneous and lagged measurements of linear and nonlinear dependence between groups of multivariate time series: frequency decomposition. ArXiv e-prints arXiv:0711.1455

Pascual-Marqui, R. D., Faber, P., Kinoshita, T., Kochi, K., Milz, P., Nishida, K., et al. (2018). A comparison of bivariate frequency domain measures of electrophysiological connectivity. bioRxiv 459503 doi:https://doi.org/10.1101/459503

Pascual-Marqui, R. D., Lehmann, D., Koukkou, M., Kochi, K., Anderer, P., Saletu, B., et al. (2011). Assessing interactions in the brain with exact low-resolution electromagnetic tomography. Phil T R Soc A 369, 3768–3784. doi:10.1098/rsta.2011.0081

Sadaghiani, S., Scheeringa, R., Lehongre, K., Morillon, B., Giraud, A.-L., D’Esposito, M., et al. (2012). Alpha-band phase synchrony is related to activity in the fronto-parietal adaptive control network. J. Neurosci. 32, 14305–14310. doi:10.1523/JNEUROSCI.1358-12.2012

Schoffelen, J. and Gross, J. (2009). Source connectivity analysis with meg and eeg. Hum Brain Mapp. 30(6), 1857–65

Soto, J. L. P., Lachaux, J.-P., Baillet, S., and Jerbi, K. (2016). A multivariate method for estimating cross-frequency neuronal interactions and correcting linear mixing in MEG data, using canonical correlations. J. Neurosci. Meth. 271, 169–181. doi:10.1016/j.jneumeth.2016.07.017

Stam, C., Nolte, G., and Daffertshofer, A. (2007). Phase lag index: assessment of functional connectivity from multi channel eeg and meg with diminished bias from common sources. Hum Brain Mapp. 28(11), 1178–93

Vinck, M., Oostenveld, R., van Wingerden, M., Battaglia, F., and Pennartz, C. M. A. (2011). An improved index of phase-synchronization for electrophysiological data in the presence of volumeconduction, noise and sample-size bias. Neuroimage 55(4), 1548–65

